# Event-related potential correlates of learning to produce novel foreign phonemes

**DOI:** 10.1101/2022.01.18.476741

**Authors:** Henry Railo, Anni Varjonen, Minna Lehtonen, Pilleriin Sikka

**Author notes:** Corresponding author Henry Railo, Assistentinkatu 7, Department of Psychology and Speech-Language Pathology, Faculty of Social Sciences, University of Turku, Finland.

## Abstract

Learning to pronounce a foreign phoneme requires an individual to acquire a motor program that enables the reproduction of the new acoustic target sound. This process is largely based on the use of auditory feedback to detect pronunciation errors to adjust vocalization. While early auditory evoked neural activity underlies automatic detection and adaptation to vocalization errors, little is known about the neural correlates of acquiring novel speech targets. To investigate the neural processes that mediate the learning of foreign phoneme pronunciation, we recorded event-related potentials (ERP) when participants (N=19) pronounced native or foreign phonemes. Behavioral results indicated that the participants’ pronunciation of the foreign phoneme improved during the experiment. Early auditory responses (N1 and P2 waves, approx. 85–290 ms after the sound onset) revealed no differences between foreign and native phonemes. In contrast, the amplitude of the fronto-centrally distributed late slow wave (LSW, 320–440 ms) was modulated by the pronunciation of the foreign phonemes, and the effect changed during the experiment, paralleling the improvement in pronunciation. These results suggest that the LSW may reflect higher-order monitoring processes that signal successful pronunciation and help learn novel phonemes.

## Introduction

When learning to pronounce foreign phonemes, the individual needs to evaluate how well the sound they produced matches the target phoneme, and in case of a mismatch, attempt to improve their phonation. Such learning may be based on fast and automatic unconscious speech control processes, but it may also depend on later processes that are associated with the conscious evaluation of phonation. However, relatively little is known about the neural mechanisms that underlie the learning of foreign phoneme production. In this study, by comparing electroencephalographic (EEG) activity evoked by self-produced and passively heard foreign and native phonemes, we investigate which auditory evoked potentials correlate with learning to pronounce foreign phonemes.

Previous research on the neural processing of foreign phonemes has focused on examining changes in auditory evoked activity elicited by passively heard sounds. Studies employing the oddball paradigm have shown that foreign phonemes elicit weaker mismatch negativity responses than native phonemes (Díaz et al., 2008; Näätänen et al., 1997; Peltola et al., 2003), indicating a poorer ability to discriminate foreign phonemes. Training improves the discrimination of phonemes, and this is reflected in the mismatch response (Tamminen et al., 2015; Tremblay et al., 1998). Phoneme discrimination training also increases the amplitude of early (N1/P2) auditory event-related potentials (ERPs), which has been interpreted as training-induced neuroplastic changes in the auditory cortex (Alain et al., 2007; Reinke et al., 2003; Saloranta et al., 2020). Additionally, studies have shown that the amplitude of a late slow wave (LSW; 300–500 ms) to passively heard phonemes correlates with phonemic learning (Alain et al., 2007; Reinke et al., 2003; Saloranta et al., 2020), but the functional role of this correlate is unclear.

Learning to discriminate the acoustic features of passively heard foreign phonemes is clearly important for learning to produce the phoneme. However, to accurately vocalize novel phonemes one needs to also learn the motor commands that enable reproduction of the desired acoustic features. The process of learning to produce novel phonemes involves tracking the auditory feedback of one’s own vocalizations to adjust future pronunciation. Studies in which speech is artificially altered (e.g., by shifting pitch) indicate that individuals can adjust their speech based on auditory feedback in just 100–150 ms (Hain et al., 2000; Houde & Jordan, 1998). This type of rapid feedback control relies on “efferent copies” of motor commands that the motor system relays to the auditory cortex to compare how well the produced speech matches the targeted motor commands (Hickok et al., 2011; Houde & Nagarajan, 2011; Tourville & Guenther, 2011). Because of this interplay between motor and sensory systems, the neural processing of passively heard and self-produced sounds differs significantly (Curio et al., 2000; Houde et al., 2002). Therefore, to understand how individuals learn novel phonemic targets, it is crucial to examine electrophysiological responses to self-produced sounds.

The electrophysiological correlate of the feedback control of self-produced speech is the suppression of auditory evoked activity. Auditory ERPs in response to self-produced sounds are suppressed in the N1 (100–200 ms), and the P2 (200–300 ms) time-windows, as compared to when the same sounds are passively heard (Behroozmand et al., 2011; Curio et al., 2000; Heinks-Maldonado et al., 2005; Houde et al., 2002; Railo et al., 2020). This phenomenon is known as the speaking-induced suppression (SIS). Using magnetoencephalography (MEG), Niziolek et al. (2013) observed that the amplitude of SIS (i.e., amplitude difference between self-produced and passively heard sounds) tracked variation in phonation: Auditory responses to phonemes that deviated from prototypical phonemes produced a decreased SIS, and the size of SIS predicted corrections to vocalization. Similarly, when speech is artificially altered, SIS decreases, suggesting that the brain detects a mismatch between feedforward motor commands and heard speech (Behroozmand et al., 2009; Behroozmand & Larson, 2011; Chang et al., 2013).

In addition to enabling online adjustments to pronunciation, such feedback control mechanism may contribute to the process of learning to correctly pronounce foreign phonemes: the individual learns to better produce the desired sound based on errors in previous vocalizations. A similar mechanism is assumed to contribute to phonemic learning in children during speech acquisition. After learning the target sound, the motor commands used to produce the sound are fine-tuned based on feedback control (Tourville & Guenther, 2011). However, we are not aware of any studies that have examined whether SIS amplitude tracks the process of learning to produce a novel foreign phoneme. Many studies have investigated sensorimotor adaptation of speech to altered auditory feedback, but such learning effects are relatively short-lived and represent changes to motor programs the individual already knows (e.g., Houde & Jordan, 1998; Lametti et al., 2018). In contrast, when learning a novel phoneme, the individual needs to learn completely novel sensorimotor speech targets.

Online adjustments to speech are typically considered rapid low-level processes, but the processes of learning to pronounce foreign phonemes may also depend on higher-order processes that take more time to unfold. SIS typically refers to effects observed in the N1 and P2 time windows, but the comparison of actively spoken and passively heard phonemes sometimes also reveals differences in later time-windows where a LSW (300–500 ms) is often observed in speech tasks. Differences in this late time-window are often neglected, likely because they are too late to reflect automatic feedback speech control, but also because there is no theoretical framework within which to interpret these findings. Whereas the classical SIS time-windows (N1, P2) likely corresponds to the automatic preconscious evaluation of self-produced speech, the later time-windows may correspond to attentive evaluation of pronunciation. Because the attentive evaluation of self-produced speech is likely to contribute to phonemic learning, and because earlier studies suggest that LSW may correlate with phoneme discrimination learning (Alain et al., 2007; Reinke et al., 2003; Saloranta et al., 2020), it is important to investigate not only the earlier time-windows (N1 and P2) but also the LSW.

Here, we recorded auditory ERPs when Finnish participants reproduced a native /ö/ phoneme or a foreign Estonian /õ/ phoneme (Speak condition) after hearing the target spoken by a native speaker. In a control condition, the participants listened to a recording of the phoneme they had just spoken (Listen condition). Comparison of the Speak and Listen conditions corresponds to the SIS, and it also allows us to test whether possible associations between pronunciation and ERP amplitudes reflect active vocalization or the passive processing of acoustic information. Assuming that participants are not able to produce the foreign phoneme as accurately as the native phoneme, we expected to observe a smaller suppression of N1 amplitudes to self-produced foreign phonemes, as compared to passively heard phonemes (i.e., reduced SIS, indicating that the auditory system detects a mismatch between produced and attempted sounds). Furthermore, assuming that participants learn to better pronounce the foreign phoneme during the experiment, we analyzed the trial-by-trial changes in ERP amplitudes in three time-windows (N1, P2, and LSW) to test whether changes in ERP amplitudes track improvements in pronunciation.

## Materials and Methods

### Participants

Twenty-one participants (students at the University of Turku) volunteered for this study. All participants were Finnish, with normal hearing and with no diagnosed learning disabilities or neurological disorders. All participants were monolingual and reported no previous experience in learning Estonian. Two participants were excluded from statistical analyses due to excessive noise in EEG. Thus, the final sample included 19 participants (range 18-35 years; 17 females, 2 males). The study was conducted according to the principles of the Declaration of Helsinki. All participants provided informed consent to participate in the study. The experiment was approved by the Ethics Committee for Human Sciences at University of Turku.

### Stimuli and Procedure

An overview of a single experimental trial is presented in Figure 1A. Participants heard a recording of the Estonian phoneme /õ/, or the Finnish phoneme /ö/, in a random order. We call this the Cue stimulus condition. The phoneme /õ/ is not part of the Finnish phonological system and was therefore unfamiliar to the subjects. Acoustically, it resembles the Finnish phoneme /ö/ (e.g., Näätänen et al., 1997). After hearing the Cue phoneme, participants attempted to repeat it as well as possible (Speak condition). After repeating the sound, they heard a playback of their produced phoneme (Listen condition). The stimuli were separated by approximately a 2–3 sec. interval. Intertrial interval was about 4 seconds. This process was repeated 50 times in a block, and the experiment consisted of five blocks (250 repetitions in total during the experiment). There was a 2–5 min break between blocks.

**Figure 1.**
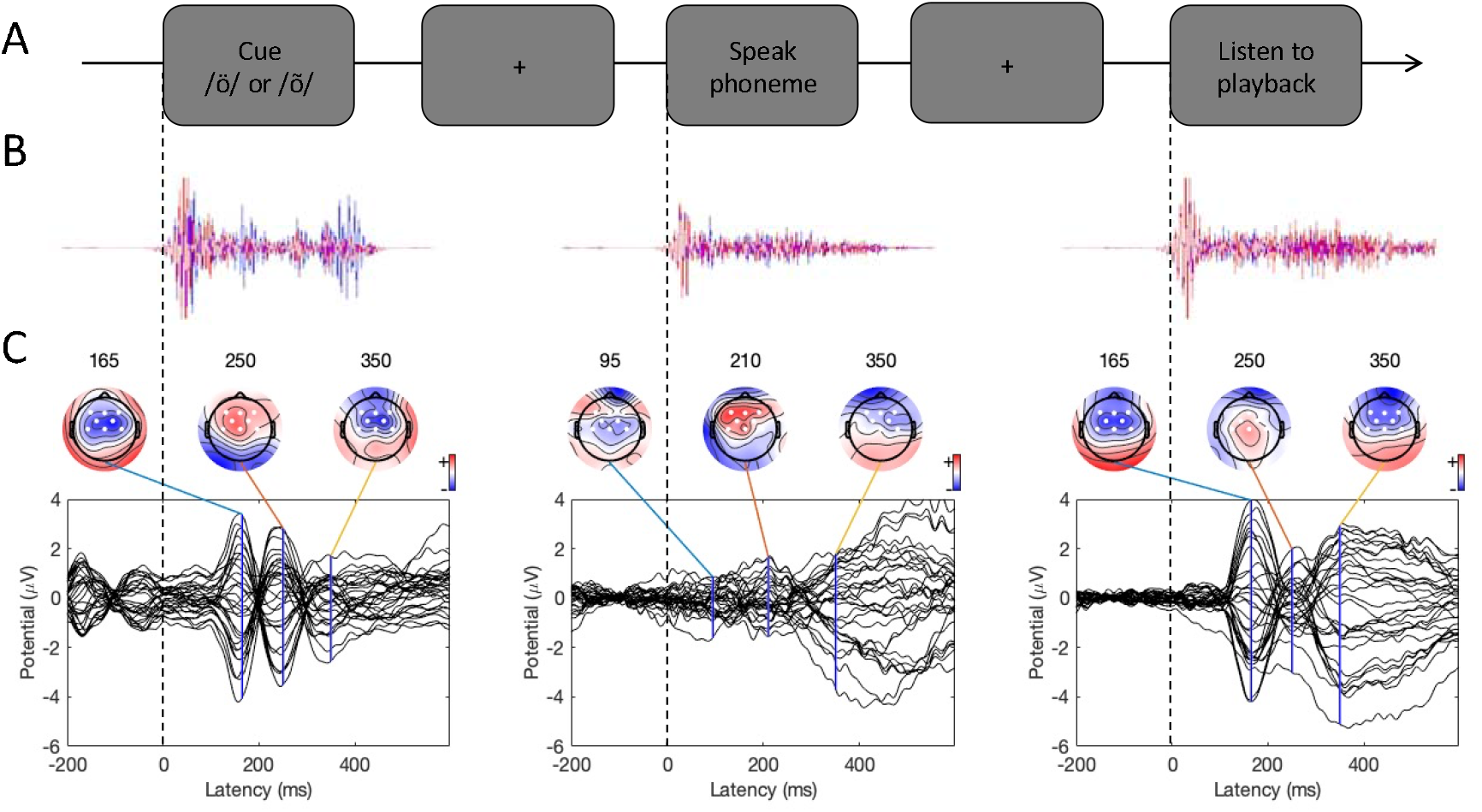
Experimental procedure and ERPs. A) An overview of a single experimental trial. B) Auditory signal recorded with the microphone (average across all participants; blue line = foreign phoneme, red = native phoneme). C) Butterfly ERP plots and scalp maps at three latencies. Scalp color map ranges from −4 (blue) to 4 μVolt (red). White electrode markers indicate the cluster of electrodes on which the statistical analysis was performed.

Cue stimuli (/ö/ and /õ/) were recorded by the same native Estonian (female) as both phonemes are part of the Estonian language. The two phonemes were approximately the same amplitude, pitch, and duration (500 ms). To examine how Finnish individuals hear the stimuli, we asked 21 participants (age range 20–50, 13 females; different participants than those who took part in the main study reported in this article) to listen to the stimuli, to identify the phoneme, and to rate how confident (scale 1–4) they were in their response. Twenty out of the 21 participants correctly identified the native phoneme as /ö/ with high confidence (M = 3.33, SD = 0.86). None of the participants correctly identified the Estonian /õ/ phoneme: Fourteen participants identified it as Finnish /ö/ (confidence M = 3.14, SD = 0.77), two participants identified the phoneme as Finnish /y/ (M = 2.0, SD = 0), two as the Russian _*bl*_ (M = 3.50, 0.70), two as the Swedish *å* (M = 4, SD = 0), and one participant believed the phoneme corresponded to the pronunciation of the German *ü* (confidence = 4). In sum, this result shows, that Finnish individuals recognize the native phoneme, but misidentify the Estonian phoneme.

The Cue and Listen stimuli were played to the participants from two TEAC LS-X8 speakers placed about 1 meter to left and right of the participant. Participants’ pronunciations were recorded using a GXT 242 Lance microphone and saved in wave file format.

### EEG Recording

EEG was recorded with 32 passive electrodes placed according to the 10-10 electrode system (EasyCap GmbH, Herrsching, Germany). Surface electromyograms (EMGs) were measured with two electrodes above and below the lips, and below and to the side of the right eye. Reference electrode was placed on the nose. Ground electrode was placed on the forehead. EEG was recorded with a NeurOne Tesla amplifier using 1.4.1.64 software (Mega Electronics Ltd., Kuopio, Finland). Sampling rate was 500 Hz. In addition, the auditory stimuli (Cue, Speak, and Listen conditions) were recorded as EEG signals using a microphone. This allowed us to accurately mark the onset times of auditory stimuli on the EEG.

### EEG Preprocessing

EEG was processed using EEGLAB v14.1.1 software (Delorme & Makeig, 2004) in Matlab 2014b. First, we high-pass filtered the microphone signals recorded with EEG at 100 Hz (to remove noise but keep the sound signal and its transient onset) and used it to add markers of the onsets of the stimuli on the continuous EEG data. The onset of auditory stimuli was determined automatically as follows. The microphone signal had to remain above a specified amplitude threshold for ten consecutive samples, and then a marker was added to the sample where the threshold was first crossed. As shown in Figure 1B, this procedure yielded accurate estimates of phoneme onset times in all three experimental conditions. After this, the microphone channels were removed from the data.

We rejected channels containing artifacts using EEGlab’s pop_rejchan function based on kurtosis, spectrum, and probability measures. Then, data were 1 Hz high-pass filtered (Klug & Gramann, 2021) using pop_eegfiltnew function, and 50 Hz line noise was reduced using the ZapLine plugin (de Cheveigné, 2020). We used Artifact Substance Reconstruction (ASR) to clean continuous EEG using a cutoff parameter at 20 (Chang et al., 2020). We then average-referenced the data and ran Independent Component Analysis (ICA; extended infomax algorithm). After the ICA, we used the DIPFIT plug-in for localizing equivalent dipole locations of the independent components. The rejection threshold was set at 100 (no dipoles were rejected) and two dipoles constrain in symmetry. We used IClabel to automatically categorize components into brain-based and various non-brain-based categories (Pion-Tonachini et al., 2019). Components with residual variance < 15%, and the probability that the component is brain based > 70%, were considered brain-based (i.e., other components were removed). After this, the removed channels were interpolated.

Next, the data was low-pass filtered at 40 Hz and cut into segments starting 200 milliseconds before stimulus onset and ending 600 milliseconds after stimulus onset. Artifactual trials were removed using the pop_jointprob function (local and global thresholds = 3). The average number of trials per participant in the Finnish Cue condition was 97 (median = 99.5, SD = 9.1) and in the Estonian Cue condition 109 (median = 112, SD = 10.1). The average number of trials in the Finnish Speak condition was 75 (median = 78, SD = 21.0) and in the Estonian Speak condition 88 (median = 99.5, SD = 28.3). In the Finnish Listen condition, the average trial number was 85 (median = 87, SD = 14.4) and in the Estonian Listen condition 95 (median = 98.5, SD = 16.4).

### Statistical Analyses

We used mixed-effects linear regression analysis to test if Condition (Speak vs. Listen) and Phoneme (Native /ö/ vs. Foreign /õ/) factors influenced ERPs at N1, P2, and LSW timewindows in single-trial data. The benefit of mixed-effects models is that analysis can be performed on single-trial data while taking into account variation between participants. The analyses were performed on a central-frontal electrode cluster (average of amplitudes across electrodes F3, Fz, F4, FC1, FC2, C3, Cz, and C4, shown in Figure 1C). The analysis was run in Matlab 2014b. The Listen condition and Native phoneme were set as reference categories in the regression models (i.e., intercept is the Native phoneme in the Listen condition). In addition, trial number was included in the model as a continuous regressor, because we were interested in examining if ERP amplitudes changed as the experiment progressed (possibly due to learning). Each condition had its own running trial number (e.g., trial number 10 in Speak/Foreign indicated trial 10 in this specific condition). The trial number regressor was z scored (i.e., intercept of the model represents responses around experiment midpoint). The model included all the three predictors and their interactions as fixed-effects regressors (i.e., Condition × Phoneme × Trial number). The model included the intercept and Speak/Listen conditions as participant-wise random effects because more complex random effect structures (based on Akaike and Bayesian information criteria, AIC/BIC) led to inferior models. Separate models were fitted for each ERP component (N1, P2, and the Slow-Wave, as described below), and the models were pruned by removing outliers.

To examine participants’ ability to pronounce the foreign phoneme, recordings of the participants’ pronunciation were rated by two native Estonians (one of the raters was coauthor P.S.). During the rating procedure, the recordings were presented in random order, one participant at a time. Participants’ pronunciation was rated on a scale from 1 to 4. Rating 1 denoted vocalizations that did *not* resemble /õ/ *at all* (e.g., the participant vocalized the phoneme /ö/). Rating 2 denoted *poor* vocalizations that had a little resemblance to /õ/, but the produced phoneme still clearly deviated from the target. Rating 3 denoted *good* vocalizations that clearly resembled /õ/ without capturing the sound perfectly. Rating 4 corresponded to an *excellent* pronunciation of /õ/ (i.e., the pronunciation resembled a native speaker’s pronunciation of the phoneme). After examining interrater correlations (Fig. 2C), the two ratings were averaged for statistical analyses. Pronunciation of native phonemes was not rated because, being part of Finnish phonetics, we assumed that participants have no difficulties in pronouncing these.

**Figure 2.**
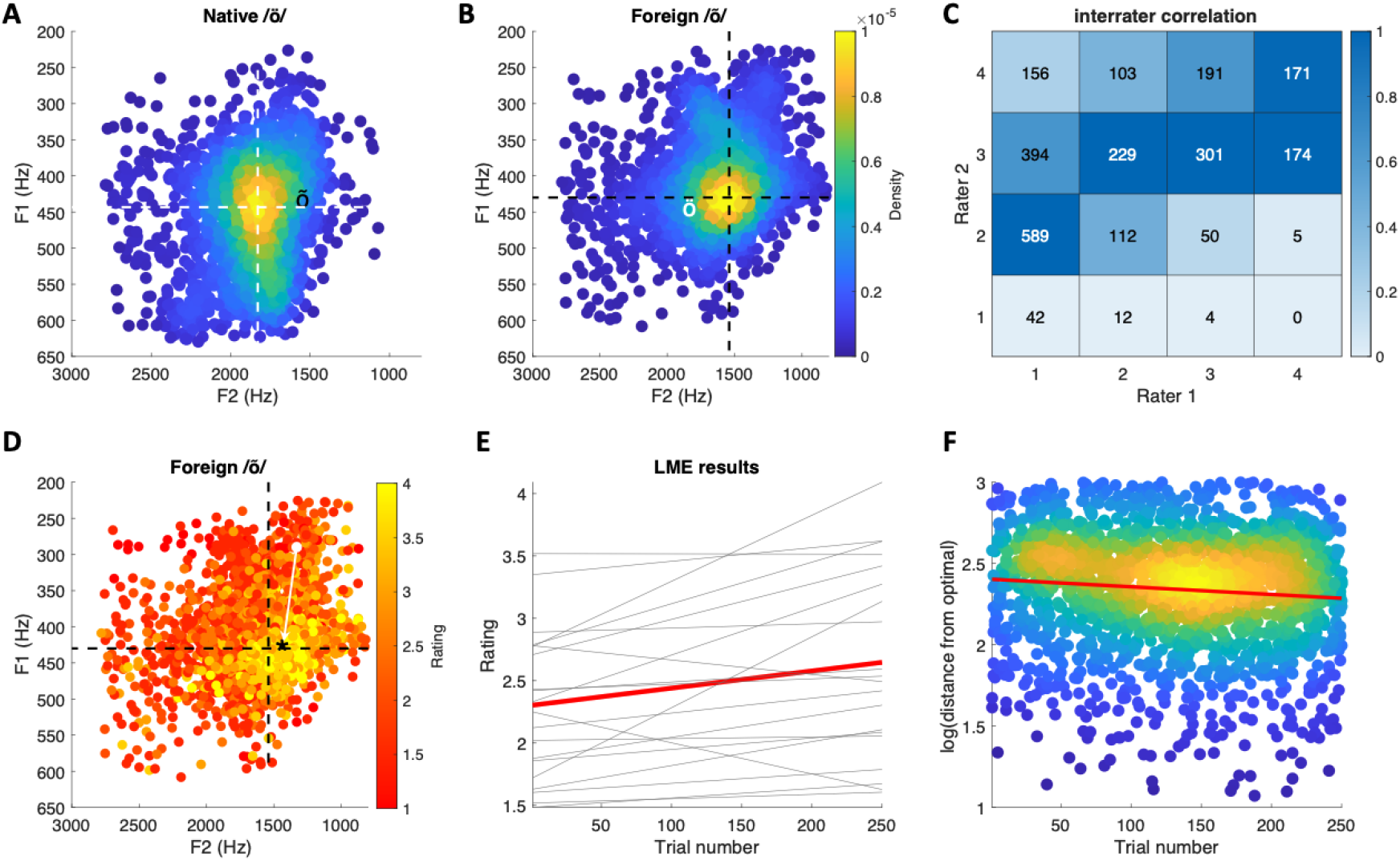
Behavioral results. A) Formant plot of single-trial estimates of native phoneme /ö/. The white dashed lines indicate the center of mass of the formant distribution. The black ‘õ’ symbol indicates the center of mass of the foreign phoneme. B) Formant plot of singletrial estimates of foreign phoneme /õ/. The black dashed lines indicate the center of mass of the formant distribution. The white ‘ö’ symbol indicates the center of mass of the native phoneme. Color code indicates the density of data points. C) Interrater correlation in phoneme ratings. Color indicates proportion of ratings (scaled per column). D) Phoneme plot of foreign /õ/ pronunciation color coded by rating. The ‘*’ symbol indicates the center of mass of vocalizations with the highest rating, and the arrow indicates the distance of a single observation from this point. E) Results of the linear mixed-effects model showing that ratings increased as the function of trials. The red line is the group average, and light lines indicate different participants’ results. F) As trial number increases, acoustic features of the foreign phoneme approach the target. The red line depicts the mixed-effects regression results (group average). Scatter density plots were made using (Nils, 2021).

The aim of the ratings was to obtain a numerical index reflecting the external validity of pronunciation. To verify that the ratings reflect acoustic parameters of the target phoneme, we examined how well the frequency of the two first formants of participants’ vocalizations predicted the (averaged) ratings. The first and second formats of the participants’ single-trial vocalizations were estimated from the wave files using linear predictive coding in Matlab.

The data, preprocessing, and analysis scripts are available at https://osf.io/wnd2j/.

## Results

### Behavioral results

Figure 2A and 2B show phoneme plots of participants’ native and foreign phoneme pronunciations, respectively. Consistent with previous reports, pronunciation of the two phonemes differ especially in F2 (e.g., Asu & Teras, 2009; Näätänen et al., 1997). The ratings of the two native Estonian speakers moderately correlated with each other (rho = .54; linearly weighed Cohen’s kappa = 0.19; Fig. 2C), but suggested that the participants had, in general, difficulties in pronouncing the foreign phoneme (mean rating = 2.46, SD = 0.81). To verify that the ratings captured systematic variation in the acoustics of the foreign phoneme, we used linear mixed-effects models to examine how well the F1 and F2 frequencies predicted the ratings. The model included F1 and F2 frequencies and their interaction as fixed-effects predictors, and the same regressors and the intercept were included as by-participant random effects variables. The results indicated that F1 frequency (t = 1.97, p = .04), F2 frequency (t = 4.06, p < .001), and their interaction (t = 2.83, p = .004) predicted roughly 67% (adjusted R^2^) of variation in the ratings.

Additionally, linear mixed-effects model indicated that, on average, ratings of participants’ pronunciation of the foreign phoneme improved as a function of trials (β = .0014, t = 2.87, p = .004; the model included random, by-participant variation in the intercept and trial number regressor). This result is visualized in Figure 2E, which shows the model results for individual participants (thin black lines) in addition to the group average (red line). Finally, to verify that the increase in ratings as a function of trials reflected true changes in the acoustic features of pronunciation, we calculated the Euclidean distance of single-trial foreign phoneme vocalizations from the “optimal” F1 and F2 formant combination (see blue arrow in Figure 2D for visualization). Linear mixed-effects model with trial number as a predictor indicated that formants approached “optimal” as a function of trial (t = −2.60; random-effect structure included by-participant intercept and trial number).

### ERP results

Butterfly plots of the ERPs in different experimental conditions are shown in Figure 1C. In both the Cue and Listen conditions, prominent N1 (negative peak at 160 ms) and P2 (positive peak at 250 ms) waves were observed. In addition, the LSW was observed around 350–500 ms after stimulus onset. As expected, the amplitude of the ERPs was suppressed in the Speak condition relative to the Listen condition, indicating SIS. In addition, the peak latency of the N1 and P2 waves were earlier, and the scalp topography somewhat different in the Speak condition (as compared to the Listen and Cue conditions).

The critical comparison between the Listen and Speak conditions is visualized in Figure 3. Scalp distributions of the difference between the Listen and Speak conditions—reflecting SIS—are shown in Figure 3B, separately for the Native and Foreign phoneme conditions. We statistically analyzed how the experimental manipulations influenced the N1 (150–200 ms), P2 (230–290 ms), and LSW (320–440 ms) amplitudes. Based on the scalp maps of prominent waves (see Figure 1C), statistical analyses were performed on the central-frontal electrode cluster (F3, Fz, F4, FC1, FC2, C3, Cz, C4).

**Figure 3.**
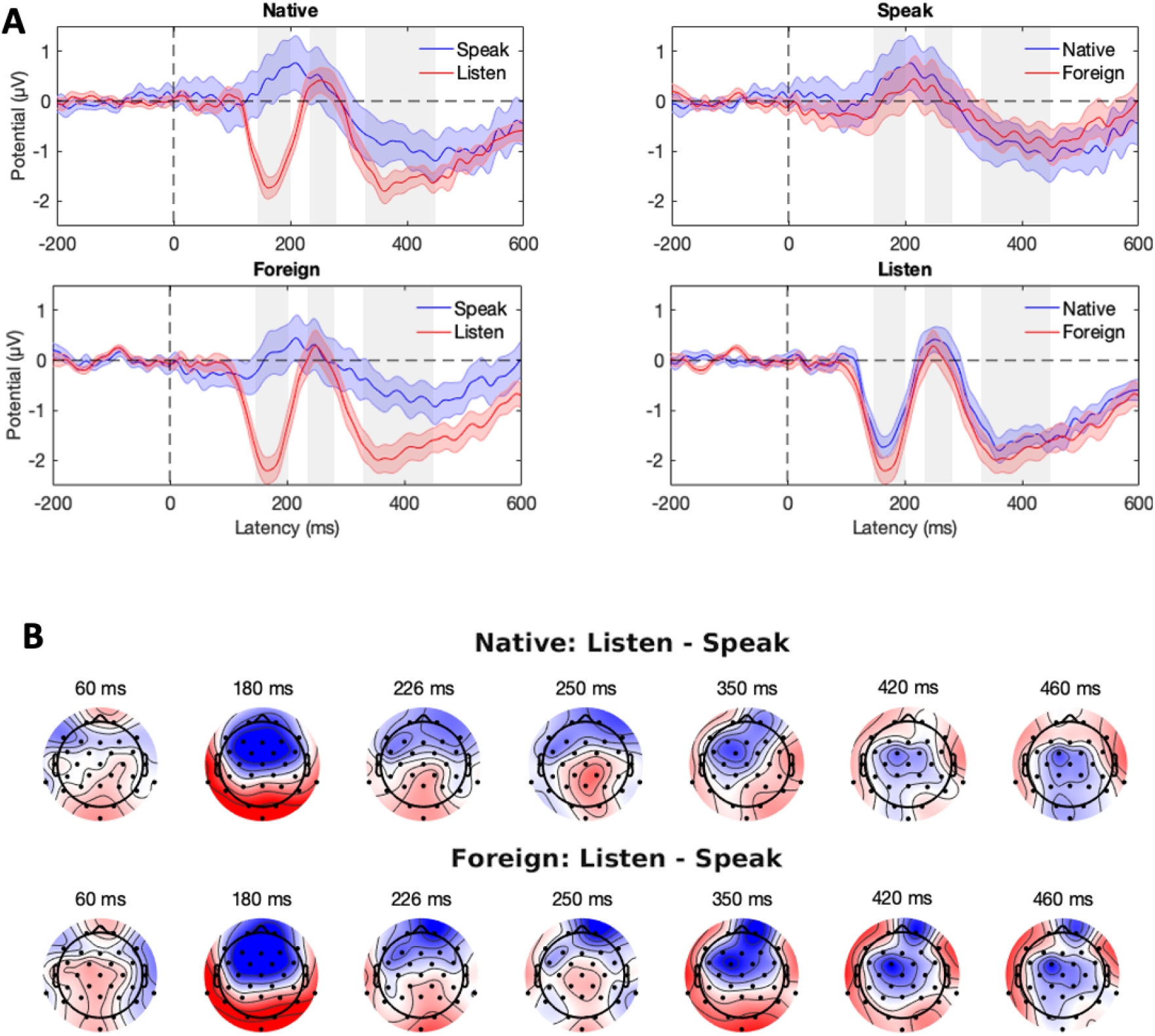
Speaking-induced suppression (SIS). A) Grand-average ERPs on the frontal central electrode cluster (F3, Fz, F4, FC1, FC2, C3, Cz, and C4). The shaded area is the standard error of the mean. The gray rectangles indicate the N1, P2, and LSW time-windows, respectively. B) Scalp maps show the SIS separately for the native and foreign phoneme conditions. Color bar ranges from −2 (blue) to 2 μV (red).

### N1 amplitudes

The results of the linear mixed-effects regression analyses on N1 amplitudes are presented in Table 1. The intercept represents the average N1 amplitude in the Listen/Native condition. The effect of Trial shows that the amplitude does not change statistically significantly (p = .17) as a function of trial. The main effect of the Speak condition indicates that, on average, amplitudes were 1.94 μV more positive in the Speak condition than in the Listen condition (p < .001), reflecting the SIS. This effect did not change statistically significantly during the experiment (Trial:Speak interaction, p = .67). N1 amplitudes to foreign phonemes were, on average, amplified by 0.38 μV (p < .001). The lack of Speak:Foreign interaction (p = .63) indicates that the SIS did not differ between the Foreign and Native conditions. Finally, the lack of a three-way Trial:Speak:Foreign interaction indicates that the SIS in the Foreign condition was not modulated by trial number (p = .88). The results of the model did not change markedly if the model was pruned by removing the (largely redundant) Trial regressor.

**Table 1.** Results of the mixed-effects regression analysis on N1 amnlitudes (df = 6157)

| Name | Estimate | SE | t | p | Lower 95%-CI | Upper 95%-CI |
| --- | --- | --- | --- | --- | --- | --- |
| Intercept | -1.54 | 0.19 | -7.98 | <.001 | -1.92 | -1.16 |
| Trial | 0.12 | 0.09 | 1.37 | 0.17 | -0.05 | 0.29 |
| Speak | 1.94 | 0.45 | 4.30 | <.001 | 1.06 | 2.83 |
| Foreign | -0.38 | 0.12 | -3.21 | <.001 | -0.62 | -0.15 |
| Trial:Speak | 0.06 | 0.13 | 0.43 | 0.67 | -0.20 | 0.31 |
| Trial:Foreign | -0.19 | 0.12 | -1.62 | 0.11 | -0.43 | 0.04 |
| Speak:Foreign | 0.09 | 0.18 | 0.49 | 0.63 | -0.26 | 0.43 |
| Trial:Speak:Foreign | -0.03 | 0.18 | -0.15 | 0.88 | -0.37 | 0.32 |

Because the latency of the N1 wave peaked earlier in the Speak than in the Listen condition, we repeated the analysis with the N1 amplitudes between 84–104 ms in the Speak condition. The overall pattern of results was similar to that reported in Table 1.

### P2 amplitudes

The results of the analyses regarding the P2 time-window are shown in Table 2. Here, the only marginally statistically significant effect is the reduced P2 amplitude in the Foreign condition (p = .05). The results did not markedly change when the model was pruned by removing the Trial regressor.

**Table 2.** Results of the mixed-effects regression analysis on P2 amplitudes (df = 6143)

| Name | Estimate | SE | t | p | Lower 95%-CI | Upper 95%-CI |
| --- | --- | --- | --- | --- | --- | --- |
| Intercept | 0.21 | 0.27 | 0.79 | 0.43 | -0.32 | 0.74 |
| Trial | 0.01 | 0.09 | 0.11 | 0.92 | -0.16 | 0.18 |
| Speak | -0.09 | 0.46 | -0.20 | 0.84 | -1.00 | 0.82 |
| Foreign | -0.23 | 0.12 | -1.94 | 0.05 | -0.46 | 0.00 |
| Trial:Speak | 0.20 | 0.13 | 1.52 | 0.13 | -0.06 | 0.45 |
| Trial:Foreign | -0.21 | 0.12 | -1.76 | 0.08 | -0.44 | 0.02 |
| Speak:Foreign | 0.12 | 0.18 | 0.67 | 0.50 | -0.23 | 0.46 |
| Trial:Speak:Foreign | 0.11 | 0.18 | 0.63 | 0.53 | -0.23 | 0.45 |

### LSW amplitudes

The results regarding the LSW time-window are shown in Table 3. As indicated by the main effect of Speak condition, the estimated SIS was 0.67 μV in the Native condition (p = .05). In the Foreign condition, the LSW amplitude for passively heard phonemes was slightly larger (0.26 μV) when compared to the Native condition (p = .04). The SIS was 0.43 μV larger in the Foreign condition than in the Native condition (Speak:Foreign, p = 0.02). Finally, the Trial:Speak:Foreign interaction (p = .01) suggests that the difference in the SIS between Foreign and Native conditions changed throughout the experiment. The Trial:Speak:Foreign interaction coefficient indicates that, as trial number increased one z unit (roughly 30 trials), the amplitude of the Speak:Foreign interaction increased by 0.49 μV.

**Table 3.**
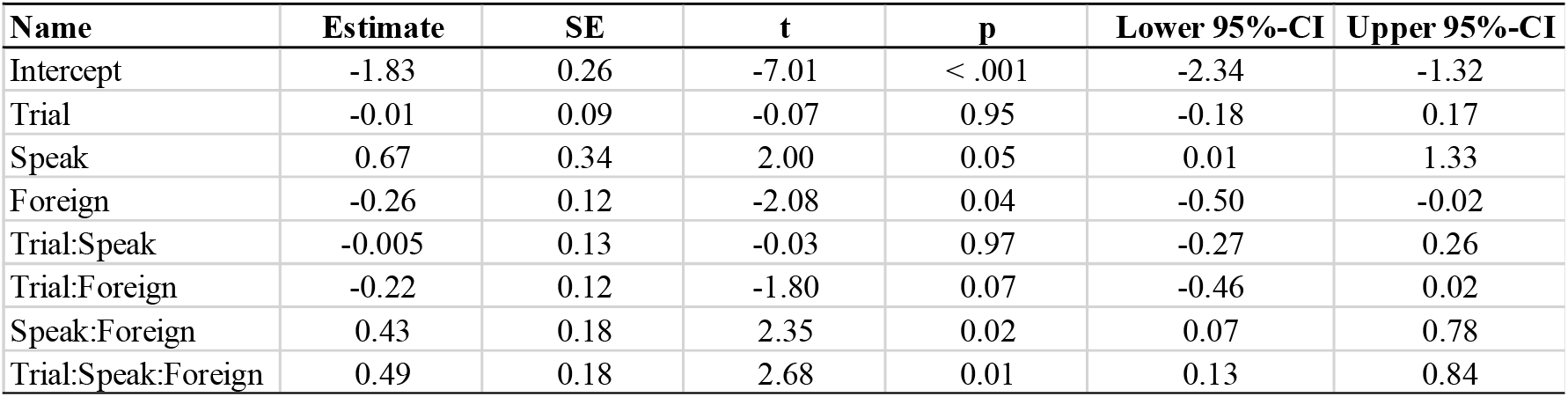
Results of the mixed-effects regression analysis on LSW amplitudes (df = 6123)

| Name | Estimate | SE | t | p | Lower 95%-CI | Upper 95%-CI |
| --- | --- | --- | --- | --- | --- | --- |
| Intercept | -1.83 | 0.26 | -7.01 | < .001 | -2.34 | -1.32 |
| Trial | -0.01 | 0.09 | -0.07 | 0.95 | -0.18 | 0.17 |
| Speak | 0.67 | 0.34 | 2.00 | 0.05 | 0.01 | 1.33 |
| Foreign | -0.26 | 0.12 | -2.08 | 0.04 | -0.50 | -0.02 |
| Trial:Speak | -0.005 | 0.13 | -0.03 | 0.97 | -0.27 | 0.26 |
| Trial:Foreign | -0.22 | 0.12 | -1.80 | 0.07 | -0.46 | 0.02 |
| Speak:Foreign | 0.43 | 0.18 | 2.35 | 0.02 | 0.07 | 0.78 |
| Trial:Speak:Foreign | 0.49 | 0.18 | 2.68 | 0.01 | 0.13 | 0.84 |

Figure 4 visualizes the modelled LWS amplitude and the resulting SIS at three different phases of the experiment. For native phonemes (blue lines), the LSW amplitude, and consequently the SIS, stayed approximately constant throughout the experiment. In contrast, the LSW evoked by the foreign phonemes (orange lines) changed across the trials: whereas in the Listen condition the LSW amplitudes became more negative, in the Speak condition they became more positive. As a result, the SIS increased throughout the experiment (from 0.28 μV to 1.91 μV).

**Figure 4.**
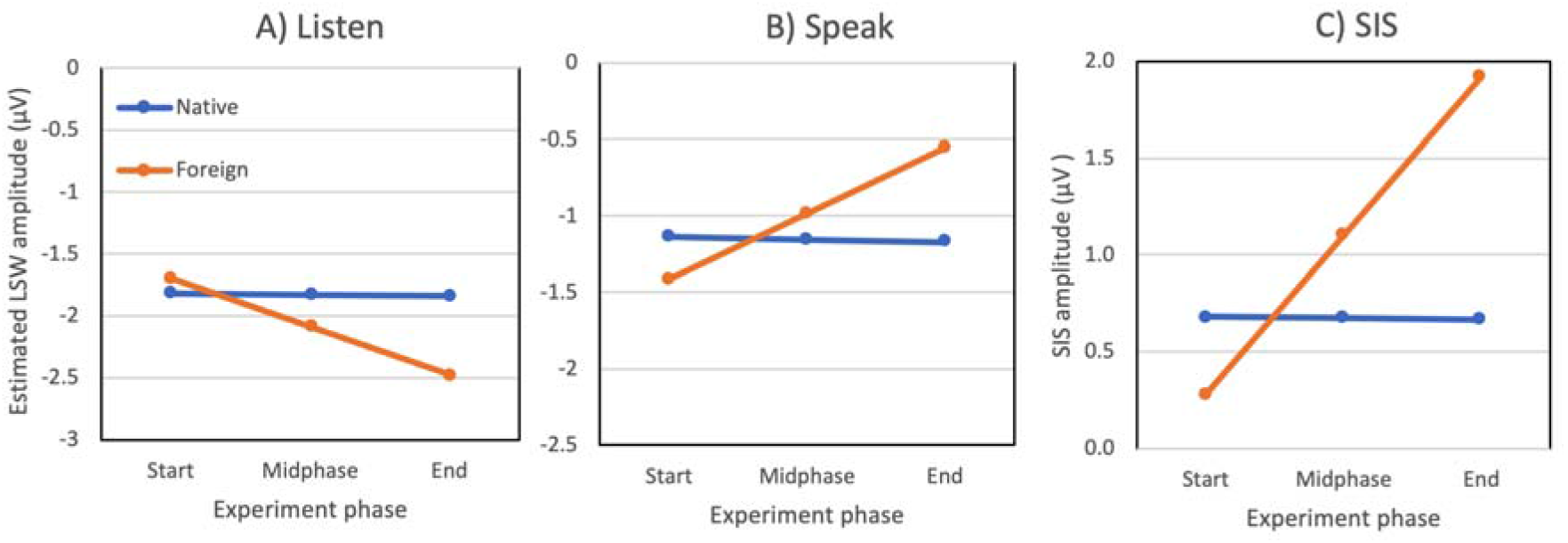
Estimated late slow wave (LSW) amplitudes in the A) Listen and B) Speak conditions, and C) the corresponding speaking-induced suppression (SIS), during different phases of the experiment. The blue line represents the Native phoneme condition and the orange line the Foreign phoneme condition. The SIS evoked by foreign phonemes increased throughout the experiment, whereas it stayed constant for native phonemes. The experiment phase refers to the Trial regressor in Table 3 (i.e., −1.7, 0, and 1.7 z-units at the Start, Midphase and End of the experiment, respectively).

### Correlation between pronunciation ratings and ERPs

The results of the analyses performed so far indicate that foreign phonemes modulate LSW amplitudes, and that this effect changes as a function of trials. To test if these changes correlate with improvements in the pronunciation of the foreign phoneme, we performed mixed-effects regression analyses to examine if differences in N1, P2, and LSW amplitudes were predicted by the ratings of foreign phoneme pronunciation accuracy (Rating). The model also included the Speak factor (Speak vs. Listen condition), and the interaction between Speak and Rating. The grand-average ERPs are presented in Figure 5A. In the N1 time-window (Figure 5B), the main effect of Rating (p = .08) and Speak:Rating interaction (p = .27) indicated that N1 amplitudes were not modulated by the accuracy of pronunciation. When the N1 amplitude in the Speak condition was calculated based on an earlier time-window (84–140 ms), the main effect of Rating was statistically significant (p = .03; Speak:Rating interaction p = .27), suggesting that higher ratings were associated with enhanced N1 amplitudes in both Speak and Listen conditions. P2 amplitudes (Figure 5C) did not change as a function of trial in the Listen condition (main effect of Rating, p = .63), but P2 was enhanced for phonemes with higher ratings (Speak:Rating, p = .014). As shown in Figure 5D, a similar pattern was observed in the LSW time-window (main effect of Rating: p = .68; Speak:Rating interaction: p = .048).

**Figure 5.**
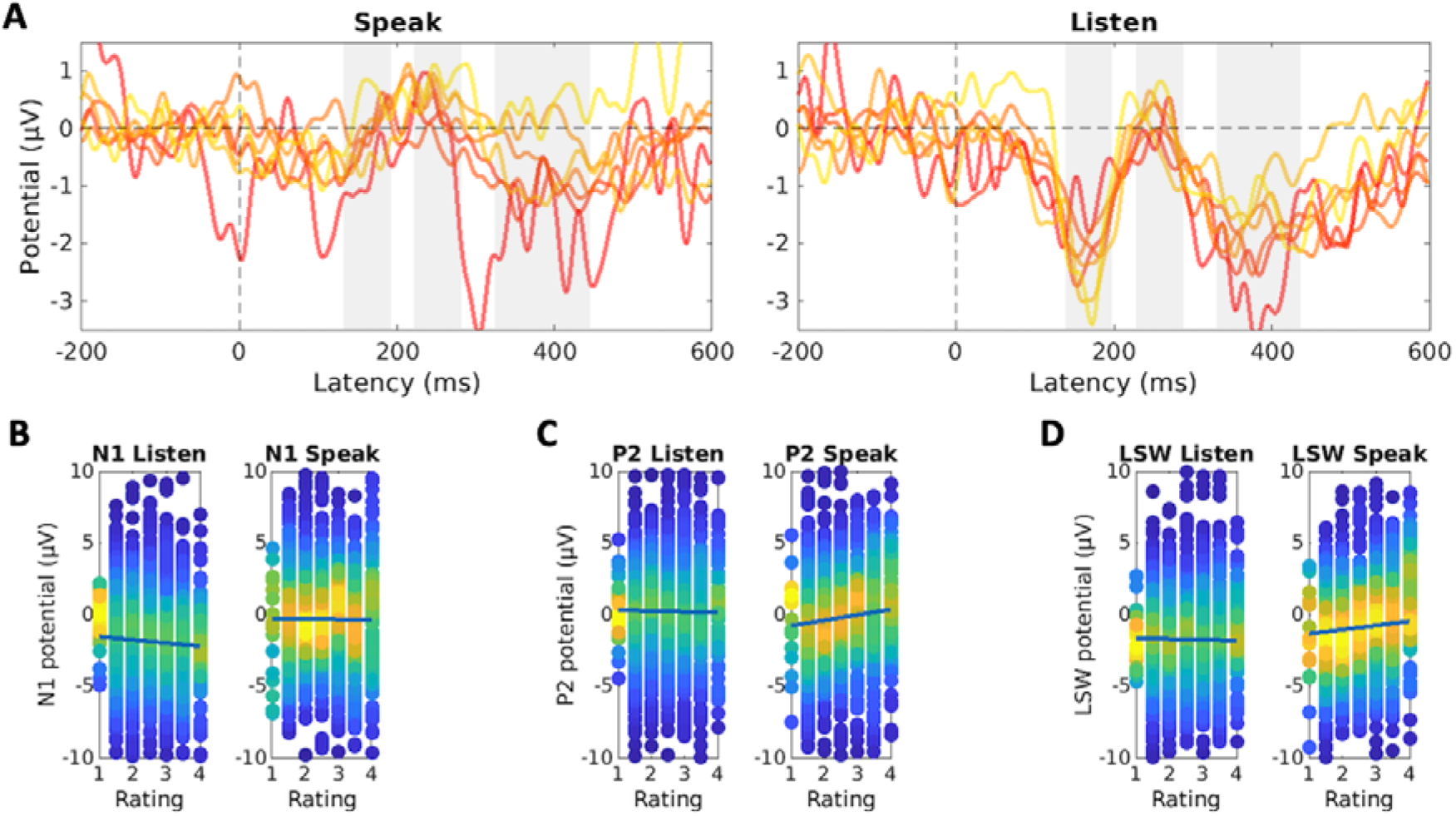
ERPs in the Foreign condition as a function of pronunciation rating. A) Grand-average ERPs for different rating categories in the Speak and Listen conditions (frontal-central electrode cluster). The gray rectangles indicate the N1, P2, and LSW time-windows. Yellow lines represent the highest ratings, red lines the lowest (see, Fig. 2D). B) N1 amplitude as a function of rating, separately for Speak and Foreign conditions. C) P2 amplitude as a function of rating, separately for Speak and Foreign conditions. D) LSW amplitude as a function of rating, separately for Speak and Foreign conditions. In panels B-D the dots indicate single-trial ERP amplitudes, and the color represents the density of the observations. The lines in panels B-D display the results of the linear mixed-effects regression model.

### Analysis of Cue sounds

We analyzed the ERPs produced by the Cue stimuli using linear mixed-effects regression models. The analysis included a factor indicating Condition (Native vs. Foreign phoneme), running (z scored) trial number, and their interaction. This analysis tests to what extent differences in the Listen and Speak conditions can be observed for stimuli not pronounced by the participants. In the N1 time-window, foreign phonemes produced stronger ERPs (β = −.25, t = −1.92, p = .054), and this effect increased through the experiment (β = −.20, t = −2.02, p = .042). Models for P2 and LSW amplitudes did not reveal any statistically significant effects.

## Discussion

When learning to pronounce a novel foreign phoneme, the individual needs to acquire novel speech targets based on auditory feedback and by trial-and-error. We examined whether a well-known marker of auditory feedback control—speaking-induced suppression (i.e., SIS) of auditory evoked activity—is modulated when participants pronounce foreign vs. native phonemes. We observed that specifically activity in a relatively late time-window (LSW, about 300 ms after vocalization onset) was modulated by the pronunciation of foreign phonemes. This effect is interesting because only LSW amplitudes evoked by *self-produced* foreign phonemes (but not passively heard foreign phonemes) changed over the course of the experiment, and this effect correlated positively with pronunciation accuracy. When participants vocalized native phonemes, the LSW remained constant throughout the experiment. Because the change in the LSW response parallels improvements in pronunciation, and the effect was specific to pronouncing foreign phonemes, we suggest this result reflects neural processes that mediate the acquisition of novel phonemes.

In line with a large body of research (Behroozmand et al., 2011; Behroozmand & Larson, 2011; Curio et al., 2000; Heinks-Maldonado et al., 2005; Houde et al., 2002; Knolle et al., 2019; Niziolek et al., 2013), we observed a strong SIS. The question arises as to why we did not observe differences between foreign and native conditions in SIS in the classical N1 and P2 time-windows. Behavioral results indicated that participants had difficulties in pronouncing the foreign /õ/. This suggests that participants did not yet have a clearly defined acoustic or motor target for the foreign phoneme, and their phonation was likely largely based on native phonemes (consistent with this, the formants of the foreign phoneme were biased towards the native /ö/ phoneme). A difference in SIS is not expected if the pronunciation of foreign phoneme relies on similar motor programs as that of native phonemes. The lack of differences in SIS between foreign and native phonemes should not therefore be taken as evidence that the mismatch between desired and produced speech does not modulate SIS. Many previous studies have shown that when vocalization does not match the target, SIS is reduced in amplitude (Behroozmand et al., 2009; Behroozmand & Larson, 2011; Chang et al., 2013; Niziolek et al., 2013). Moreover, the present findings showed that better accuracy in pronouncing foreign phonemes correlated positively with P2 amplitudes in the Speak condition. This indicates that better foreign pronunciation was associated with reduced SIS in the P2 time window.

The fact that we did not find any differences in SIS in the early time windows suggests that learning to better pronounce the foreign phoneme was not strongly associated with low-level auditory feedback control mechanisms. Instead, the change in pronunciation accuracy correlated with LSW amplitudes. When the participants passively listened to the foreign phoneme, the LSW amplitude became more negative across the experiment. This result parallels the findings reported by Alain et al. (2007) and Reinke et al. (2003) who observed that phoneme discrimination training was associated with a modulated LSW. But when the participants in the present study actively pronounced the foreign phoneme, the LSW amplitude became more positive across trials. Consequently, the SIS evoked by foreign phonemes increased as a function of trials. Control analyses of the Cue sounds (/ö/ and /õ/ phonemes spoken by a native speaker) indicated that the difference in the LSW amplitudes cannot be attributed to acoustic differences between the /ö/ and /õ/ phonemes.

What cognitive or behavioral processes does the change in the LSW amplitude to self-produced foreign phonemes reflect? Firstly, because the pronunciation of foreign phonemes improved across the experiment, the LSW amplitude does not appear to reflect fatigue or inattentiveness. Moreover, if the change in LSW reflects fatigue, LSW should be negatively associated with pronunciation accuracy, which is the opposite of what we observed. Secondly, it could be suggested that the LSW evoked by self-produced phonemes is an error signal, indicating a mismatch between the produced and attempted vocalization. However, if LSW reflects an error signal, it should correlate negatively with pronunciation accuracy. Instead, in the present study, enhanced amplitude of SIS in the LSW time-window indicated improved pronunciation. These findings suggest that the LSW may reflect processing of successful vocalizations, which enables the participant to notice and adjust future vocalizations.

The positive amplitude shift in LSW coincides with the timing and topography of the P3a wave, which reflects attentional capture of salient or motivationally relevant stimuli (Knolle et al., 2019; Polich, 2007). The P3a wave is elicited by sensory stimuli that require a behavioral response form the participant (e.g., participant needs to classify a stimulus and respond using a button press) (Pitts et al., 2012; Scheerer & Jones, 2018), which likely explains why a P3a wave was not observed in the present study. Although we did not observe a P3a wave per se, the neurocognitive basis of the positive shift in the LSW amplitude may be similar to the mechanisms producing the P3a.

Consciously noticing successful vocalizations is likely a key part of learning to pronounce foreign phonemes because the process requires identifying new acoustic speech targets; it is difficult to improve one’s vocalization of foreign phonemes unless one consciously knows what to aim for. Noticing successful pronunciations could also be interpreted as motivationally relevant reinforcement signals that drive learning. Like the LSW effect we observed, correctly performed actions and feedback after a successful performance are associated with a positive amplitude shift in frontal central locations (called the “reward positivity”), often interpreted as a correlate of reinforcement signals (Carlson et al., 2011; Glazer et al., 2018; Holroyd & Coles, 2002; Hoy et al., 2021; Ullsperger et al., 2014). Consistent with this, individuals learn better from feedback following successful trials (rather than errors) (Arbel et al., 2013; Chiviacowsky & Wulf, 2007). Although, in the present study, the participants were not provided direct feedback on the success of their pronunciation, the playback of their own pronunciation likely functioned as type of feedback signal (Parrell, 2021). Reinforcement learning likely plays a role in learning to produce foreign phonemes but the topic remains little studied (Parrell, 2021).

Based on the similarities to P3a and reward positivity, we suggest that the positive shift in the LSW amplitude to self-produced foreign phonemes may reflect motivational, saliency, or reinforcement signals. These enable the individual to notice successful pronunciations, which then translates to improved pronunciation during the experiment. By successful pronunciation we do not simply refer to the absence of vocalization errors, but to pronunciation that finds its target *surprisingly* well. These successful pronunciations could be “planned” (i.e., in the sense that this was what the individual attempted to do), but they could also be due to pronunciation errors that, by coincidence, match the target sound better than intended.

Assuming that the sources that contribute to P3a also contribute to LSW, the positive shift in the LSW amplitude could also reflect participants’ conscious confidence in their performance. Frömer et al. (2021) showed that P3a amplitudes correlated with participants’ confidence on the success of their actions. The authors suggest that confidence approximates the amount of noise in the efference copy signals (i.e., their precision), which together with the efference copy and feedback, allow the individuals to make inferences about the success of their action. Their results showed that participants with more accurately “calibrated” confidence (i.e., more accurate estimate of noise in efference copies) learned better (Frömer et al., 2021). This suggests that LSW could reflect higher-order monitoring mechanisms: Whereas SIS in the N1 time-window may reflect automatic “online” corrections based on the efference copy (Hickok et al., 2011; Houde & Nagarajan, 2011; Tourville & Guenther, 2011), the LSW may reflect monitoring processes that integrate multiple sources of information (e.g., auditory feedback on one’s pronunciation, predictions based on the efference copy, and the individual’s confidence in the accuracy of these predictions).

In conclusion, our results show that the ability to correctly pronounce a novel foreign phoneme correlates with the amplitude of a late slow ERP (320–440 ms after vocalization). Activity during this time-window was differently modulated when participants pronounced foreign (as compared to native) phonemes, and the effect changed during the experiment, paralleling improvements in pronunciation. We propose that activity in the LSW timewindow reflects high-level performance monitoring processes that signal successful pronunciations and help learn motor commands that produce novel acoustic speech targets. If our interpretation is correct, we expect that the effect in the LSW time-window (difference between foreign and native conditions) disappears once the individual has successfully learned to produce the novel phoneme (i.e., there is little room, or need for further improvement), and the pronunciation of the foreign phoneme has automatized. A better understanding of the processes that mediate the acquisition of novel phonemes may, in the future, helps shed light on various speech-related phenomena, such as the neural processes underlying native language acquisition, second language learning, and rehabilitation of speech deficits.

## Acknowledgements

We thank Ita Puusepp for help with the ratings. M.L. was partly supported by the Research Council of Norway through its Centers of Excellence funding scheme (project number 223265). P.S. was supported by a research grant from the Alfred Kordelin Foundation, and by a research grant from the Emil Aaltonen Foundation. We thank Teemu Laine for help with the experimental set up and equipment.

## Notes

### Competing Interest Statement

The authors have declared no competing interest.

### Summary of Updates

Accepted manuscript version. Minor modifications to previous versions.

## References

Alain, C., Snyder, J. S., He, Y., & Reinke, K. S. (2007). Changes in auditory cortex parallel rapid perceptual learning. Cerebral Cortex (New York, N.Y.□: 1991), 17(5), 1074–1084. https://doi.org/10.1093/CERCOR/BHL018

Arbel, Y., Goforth, K., & Donchin, E. (2013). The Good, the Bad, or the Useful? The Examination of the Relationship between the Feedback-related Negativity (FRN) and Longterm Learning Outcomes. Journal of Cognitive Neuroscience, 25(8), 1249–1260. https://doi.org/10.1162/JOCN_A_00385

Asu, E. L., & Teras, P. (2009). Estonian. Journal of the International Phonetic Association, 39(3), 367–372. https://doi.org/10.1017/S002510030999017X

Behroozmand, R., Karvelis, L., Liu, H., & Larson, C. R. (2009). Vocalization-induced enhancement of the auditory cortex responsiveness during voice F0 feedback perturbation. Clinical Neurophysiology. https://doi.org/10.1016/j.clinph.2009.04.022

Behroozmand, R., & Larson, C. R. (2011). Error-dependent modulation of speech-induced auditory suppression for pitch-shifted voice feedback. BMC Neuroscience. https://doi.org/10.1186/1471-2202-12-54

Behroozmand, R., Liu, H., & Larson, C. R. (2011). Time-dependent neural processing of auditory feedback during voice pitch error detection. Journal of Cognitive Neuroscience. https://doi.org/10.1162/jocn.2010.21447

Carlson, J. M., Foti, D., Mujica-Parodi, L. R., Harmon-Jones, E., & Hajcak, G. (2011). Ventral striatal and medial prefrontal BOLD activation is correlated with reward-related electrocortical activity: A combined ERP and fMRI study. NeuroImage, 57(4), 1608–1616. https://doi.org/10.1016/J.NEUROIMAGE.2011.05.037

Chang, C. Y., Hsu, S. H., Pion-Tonachini, L., & Jung, T. P. (2020). Evaluation of Artifact Subspace Reconstruction for Automatic Artifact Components Removal in Multi-Channel EEG Recordings. IEEE Transactions on Biomedical Engineering, 67(4), 1114–1121. https://doi.org/10.1109/TBME.2019.2930186

Chang, E. F., Niziolek, C. A., Knight, R. T., Nagarajan, S. S., & Houde, J. F. (2013). Human cortical sensorimotor network underlying feedback control of vocal pitch. Proceedings of the National Academy of Sciences of the United States of America. https://doi.org/10.1073/pnas.1216827110

Chiviacowsky, S., & Wulf, G. (2007). Feedback after good trials enhances learning. Research Quarterly for Exercise and Sport, 78(2), 40–47. https://doi.org/10.1080/02701367.2007.10599402

Curio, G., Neuloh, G., Numminen, J., Jousmäki, V., & Hari, R. (2000). Speaking modifies voice-evoked activity in the human auditory cortex. Human Brain Mapping. https://doi.org/10.1002/(SICI)1097-0193(200004)9:4<183::AID-HBM1>3.0.CO;2-Z

de Cheveigné, A. (2020). ZapLine: A simple and effective method to remove power line artifacts. NeuroImage, 207. https://doi.org/10.1016/J.NEUROIMAGE.2019.116356

Delorme, A., & Makeig, S. (2004). EEGLAB: An open source toolbox for analysis of single-trial EEG dynamics including independent component analysis. Journal of Neuroscience Methods. https://doi.org/10.1016/j.jneumeth.2003.10.009

Díaz, B., Baus, C., Escera, C., Costa, A., & Sebastián-Gallés, N. (2008). Brain potentials to native phoneme discrimination reveal the origin of individual differences in learning the sounds of a second language. Proceedings of the National Academy of Sciences, 105(42), 16083–16088. https://doi.org/10.1073/PNAS.0805022105

Frömer, R., Nassar, M. R., Bruckner, R., Stürmer, B., Sommer, W., & Yeung, N. (2021). Response-based outcome predictions and confidence regulate feedback processing and learning. ELife, 10. https://doi.org/10.7554/ELIFE.62825

Glazer, J. E., Kelley, N. J., Pornpattananangkul, N., Mittal, V. A., & Nusslock, R. (2018). Beyond the FRN: Broadening the time-course of EEG and ERP components implicated in reward processing. International Journal of Psychophysiology, 132, 184–202. https://doi.org/10.1016/J.IJPSYCHO.2018.02.002

Hain, T. C., Burnett, T. A., Kiran, S., Larson, C. R., Singh, S., & Kenney, M. K. (2000). Instructing subjects to make a voluntary response reveals the presence of two components to the audio-vocal reflex. Experimental Brain Research, 130(2), 133–141. https://doi.org/10.1007/S002219900237

Heinks-Maldonado, T. H., Mathalon, D. H., Gray, M., & Ford, J. M. (2005). Fine-tuning of auditory cortex during speech production. Psychophysiology. https://doi.org/10.1111/j.1469-8986.2005.00272.x

Hickok, G., Houde, J., & Rong, F. (2011). Sensorimotor Integration in Speech Processing: Computational Basis and Neural Organization. Neuron, 69(3), 407–422. https://doi.org/10.1016/J.NEURON.2011.01.019

Holroyd, C. B., & Coles, M. G. H. (2002). The neural basis of human error processing: reinforcement learning, dopamine, and the error-related negativity. Psychological Review, 109(4), 679–709. https://doi.org/10.1037/0033-295X.109.4.679

Houde, J. F., & Jordan, M. I. (1998). Sensorimotor adaptation in speech production. Science. https://doi.org/10.1126/science.279.5354.1213

Houde, J. F., & Nagarajan, S. S. (2011). Speech production as state feedback control. In Frontiers in Human Neuroscience. https://doi.org/10.3389/fnhum.2011.00082

Houde, J. F., Nagarajan, S. S., Sekihara, K., & Merzenich, M. M. (2002). Modulation of the auditory cortex during speech: An MEG study. Journal of Cognitive Neuroscience. https://doi.org/10.1162/089892902760807140

Hoy, C. W., Steiner, S. C., & Knight, R. T. (2021). Single-trial modeling separates multiple overlapping prediction errors during reward processing in human EEG. Communications Biology 2021 4:1, 4(1), 1–17. https://doi.org/10.1038/s42003-021-02426-1

Klug, M., & Gramann, K. (2021). Identifying key factors for improving ICA-based decomposition of EEG data in mobile and stationary experiments. European Journal of Neuroscience, 54(12), 8406–8420. https://doi.org/10.1111/EJN.14992

Knolle, F., Schwartze, M., Schröger, E., & Kotz, S. A. (2019). Auditory Predictions and Prediction Errors in Response to Self-Initiated Vowels. Frontiers in Neuroscience, 13, 1146. https://doi.org/10.3389/FNINS.2019.01146/BIBTEX

Lametti, D. R., Smith, H. J., Watkins, K. E., & Shiller, D. M. (2018). Robust Sensorimotor Learning during Variable Sentence-Level Speech. Current Biology, 28(19), 3106–3113.e2. https://doi.org/10.1016/J.CUB.2018.07.030

Näätänen, R., Lehtokoski, A., Lennes, M., Cheour, M., Huotilainen, M., Iivonen, A., Vainio, M., Alku, P., Ilmoniemi, R. J., Luuk, A., Allik, J., Sinkkonen, J., & Alho, K. (1997). Language-specific phoneme representations revealed by electric and magnetic brain responses. Nature, 385(6615), 432–434. https://doi.org/10.1038/385432A0

Nils. (2021). Scatter Plot colored by Kernel Density Estimate (https://se.mathworks.com/matlabcentral/fileexchange/65728-scatter-plot-colored-by-kernel-density-estimate). Matlab.

Niziolek, C. A., Nagarajan, S. S., & Houde, J. F. (2013). What does motor efference copy represent? evidence from speech production. Journal of Neuroscience. https://doi.org/10.1523/JNEUROSCI.2137-13.2013

Parrell, B. (2021). A Potential Role for Reinforcement Learning in Speech Production. Journal of Cognitive Neuroscience, 33(8), 1470–1486. https://doi.org/10.1162/JOCN_A_01742

Peltola, M. S., Kujala, T., Tuomainen, J., Ek, M., Aaltonen, O., & Näätänen, R. (2003). Native and foreign vowel discrimination as indexed by the mismatch negativity (MMN) response. Neuroscience Letters, 352(1), 25–28. https://doi.org/10.1016/J.NEULET.2003.08.013

Pion-Tonachini, L., Kreutz-Delgado, K., & Makeig, S. (2019). ICLabel: An automated electroencephalographic independent component classifier, dataset, and website. Neuroimage, 198, 181–197. https://doi.org/10.1016/J.NEUROIMAGE.2019.05.026

Pitts, M. A., Martínez, A., & Hillyard, S. A. (2012). Visual processing of contour patterns under conditions of inattentional blindness. Journal of Cognitive Neuroscience. https://doi.org/10.1162/jocn_a_00111

Polich, J. (2007). Updating P300: An integrative theory of P3a and P3b. In Clinical Neurophysiology. https://doi.org/10.1016/j.clinph.2007.04.019

Railo, H., Nokelainen, N., Savolainen, S., & Kaasinen, V. (2020). Deficits in monitoring self-produced speech in Parkinson’s disease. Clinical Neurophysiology. https://doi.org/10.1016/j.clinph.2020.05.038

Reinke, K. S., He, Y., Wang, C., & Alain, C. (2003). Perceptual learning modulates sensory evoked response during vowel segregation. Brain Research. Cognitive Brain Research, 17(3), 781–791. https://doi.org/10.1016/S0926-6410(03)00202-7

Saloranta, A., Alku, P., & Peltola, M. S. (2020). Listen-and-repeat training improves perception of second language vowel duration: Evidence from mismatch negativity (MMN) and N1 responses and behavioral discrimination. International Journal of Psychophysiology □: Official Journal of the International Organization of Psychophysiology, 147, 72–82. https://doi.org/10.1016/J.IJPSYCHO.2019.11.005

Scheerer, N. E., & Jones, J. A. (2018). The role of auditory feedback at vocalization onset and mid-utterance. Frontiers in Psychology. https://doi.org/10.3389/fpsyg.2018.02019

Tamminen, H., Peltola, M. S., Kujala, T., & Näätänen, R. (2015). Phonetic training and non-native speech perception--New memory traces evolve in just three days as indexed by the mismatch negativity (MMN) and behavioural measures. International Journal of Psychophysiology □: Official Journal of the International Organization of Psychophysiology, 97(1), 23–29. https://doi.org/10.1016/J.IJPSYCHO.2015.04.020

Tourville, J. A., & Guenther, F. H. (2011). The DIVA model: A neural theory of speech acquisition and production. Language and Cognitive Processes, 26(7), 952. https://doi.org/10.1080/01690960903498424

Tremblay, K., Kraus, N., & McGee, T. (1998). The time course of auditory perceptual learning: neurophysiological changes during speech-sound training. Neuroreport, 9(16), 3557–3560. https://doi.org/10.1097/00001756-199811160-00003

Ullsperger, M., Fischer, A. G., Nigbur, R., & Endrass, T. (2014). Neural mechanisms and temporal dynamics of performance monitoring. Trends in Cognitive Sciences, 18(5), 259–267. https://doi.org/10.1016/J.TICS.2014.02.009

